# Quaternion Spectral Fingerprinting of DNA: GPU-Accelerated Multi-Channel Fourier Analysis for Alignment-Free Genomics

**DOI:** 10.64898/2026.04.03.716441

**Authors:** Mohamed Amine Bergach

## Abstract

Spectral methods for DNA sequence analysis—treating genomic data as a discrete signal and computing its Fourier transform—were proposed over three decades ago but remained impractical for whole-genome analysis due to computational cost. We present a quaternion Fourier transform framework that encodes DNA as a quaternion-valued signal *q*[*n*] ∈ {1, **i, j, k**} mapping to the four nucleotides *{A, T, G, C}*, and prove that the full quaternion spectrum is computable from exactly two standard complex FFTs: *Q*(*k*) = *Z*_1_(*k*) + *Z*_2_(*N*−*k*) · **j**, where *Z*_1_ = FFT(*u*_*A*_ +*i* · *u*_*T*_ ) and *Z*_2_ = FFT(*u*_*G*_ +*i* · *u*_*C*_). We establish that the resulting spectral fingerprint *F*(*k*) = (|*Z*_1_(*k*)|^2^,|*Z*_2_(*k*)|^2^) is invariant under both cyclic shift and reverse complement— the two fundamental symmetries of double-stranded DNA.

Building on this theoretical foundation, we develop three computational tools: (i) a 4×4 Hermitian cross-spectral matrix with inter-channel coherence analysis, (ii) a genome spectrogram via sliding-window short-time Fourier transform, and (iii) an alignment-free spectral variant detection algorithm with *O*(*N* log *N*) complexity. Applying Welch’s cross-spectral coherence analysis to *E. coli* K-12, we discover that the DNA helical repeat (∼11 bp) is invisible to the standard power spectrum but clearly detected through the cross-spectral matrix condition number (*κ* = 6.5), demonstrating that multi-channel analysis reveals structural periodicities that single-channel methods miss. Phase spectrum analysis recovers the characteristic nucleotide ordering within codons (*A* → *T* → *G* → *C*), while three distinct frequency regimes of inter-nucleotide coupling emerge: complementary-dominated (long-range), purine/pyrimidine-dominated (structural), and codon-position-dominated (coding).

Cross-species validation on 18 genomes spanning all three domains of life—Bacteria (5), Archaea (3), and Eukarya (10)—with GC content from 19.6% (*P. falciparum*) to 69.5% (*T. thermophilus*) confirms the universality of these findings. The helical repeat is detected via cross-spectral coherence in 18/18 organisms (100%). All 10 eukaryotes show A-T dominance at the helical repeat—a spectral signature of nucleosome wrapping absent from prokaryotes. Non-complementary pairs (A-C, T-G) dominate the coding frequency in 17/18 organisms.

Validation on human chromosome 21 (46.7 Mb, processed in 5.0 s on Apple M1) reveals eukaryote-specific spectral signatures—nucleosome positioning at 10.67 bp, nucleosome spacing at 170.7 bp, and Alu repeat dominance at 341 bp—absent from prokaryotic spectra. A proof-of-concept spectral variant detection experiment achieves 100% read-matching accuracy (100/100 reads) and statistically significant discrimination of SNPs from sequencing errors (*t* = 14.80, *p* < 0.001, Cohen’s *d* = 1.64), scaling to *d* = 8.96 at 30× coverage. The full human genome can be spectrally analyzed in approximately 3–4 seconds on an M1 GPU and under 1 second on M4 Max, enabling interactive spectral genomics on commodity hardware.

**Availability:** Source code, Metal kernels, and benchmarks are available at https://github.com/aminems/AppleSiliconFFT under the MIT license.

## 1 Introduction

The idea of treating DNA sequences as digital signals and analyzing them in the frequency domain dates to the early 1990s, when Voss [**?**] demonstrated long-range fractal correlations in nucleotide sequences and Peng et al. [**?**] discovered 1*/f* noise patterns in genomic data. The most practically significant spectral feature—the period-3 signature of protein-coding regions arising from the triplet codon structure—was first noted by Fickett [**?**] and later formalized through Fourier analysis by Tiwari et al. [**?**] and Anastassiou [**?**].

The recognition that DNA indicator sequences form a multivariate categorical time series led Stoffer, Tyler and McDougall [**?**] to introduce the *spectral envelope*—the largest eigenvalue of the cross-spectral density matrix of the indicator sequences—as an optimal scaling for periodicity detection. Stoffer and Tyler [**?**] later extended this to cross-spectral coherence between two categorical sequences for sequence matching. Herzel, Weiss and Trifonov [**?**] identified purine-purine and pyrimidine-pyrimidine coupling at the helical repeat (∼10–11 bp) through dinucleotide cross-correlation analysis in the time domain. Brodzik [**?**] mapped nucleotides to pure quaternions for a periodicity transform, though not a full quaternion DFT.

Despite these foundations, two gaps remained: (i) only the spectral envelope (largest eigenvalue) was typically extracted, discarding the full pairwise coherence structure; and (ii) no systematic cross-species analysis of the coherence landscape had been performed. This paper addresses both gaps while also tackling the computational barrier that limited prior work to individual genomes or short segments.

This paper extends the spectral envelope framework to a full pairwise coherence analysis within a quaternion algebraic structure, and validates the results across the tree of life. Our contributions are:

1. A *quaternion Fourier transform* framework that encodes DNA as a quaternion-valued signal *q*[*n*] ∈ *{*1, **i, j, k**}, connecting the four Voss indicator channels to the algebraic structure of the quaternion DFT [**?**] with formally proven spectral symmetries (Section **??**).
2. A proof that the full quaternion spectrum is computable from exactly *two* standard complex FFTs via the symplectic decomposition, enabling direct use of GPU-optimized FFT libraries (Section **??**).
3. A proof of spectral invariance under reverse complement—to our knowledge, the first formal strand-agnostic guarantee for power-spectral DNA analysis (Section **??**).
4. Extension of the spectral envelope [**?**] to the *full* pairwise coherence structure: all six magnitude-squared coherences, phase spectra, spectral entropy, and the condition number *κ*(*k*) = *λ*_max_*/λ*_min_ at each frequency, using Welch’s cross-spectral estimation [**?**] (Section **??**).
5. Cross-species validation on 18 genomes spanning Bacteria (5), Archaea (3), and Eukarya (10) with GC content from 19.6% to 69.5%, establishing the universality of three frequency regimes of inter-nucleotide coupling (Section **??**).
6. An alignment-free spectral variant detection algorithm with *O*(*N* log *N*) complexity, with proof-of-concept achieving SNP-error discrimination at *p <* 0.001 (Section **??**).
7. GPU implementation achieving whole-genome spectral analysis in seconds on commodity Apple Silicon hardware at 138 GFLOPS (Section **??**).

**Table 1:** Comparison of DNA numerical encoding schemes. “Cross-ch.” indicates the level of inter-nucleotide spectral information retained. The spectral envelope [**?**] extracts the dominant eigenvalue; our quaternion encoding extracts all six pairwise coherences.

| Encoding | Mapping | Ch. | FFTs | Cross-ch. |
| --- | --- | --- | --- | --- |
| Voss indicator [?] | $u_X[n] \in \{0, 1\}$ | 4 | 4 real | No |
| Purine/Pyrimidine | A,G→1; T,C→-1 | 1 | 1 real | N/A |
| EIIP [?] | Base→potential | 1 | 1 real | N/A |
| Complex [?] | A→1+i, etc. | 1 | 1 cplx | Partial |
| Spectral envelope [?] | $u_X[n] \in \{0, 1\}$ | 4 | 4 real | $\lambda_{\max}$ only |
| **Quaternion (ours)** | $q[n] \in \{1, \mathbf{i}, \mathbf{j}, \mathbf{k}\}$ | **1** | **2 cplx** | **Full (6 pairs)** |

The radar-to-genomics connection is not merely metaphorical. Our FFT kernels were originally developed for Synthetic Aperture Radar (SAR) processing [**?, ?**], and the signal processing techniques we apply to DNA—matched filtering, coherent integration, spectral fingerprinting—are direct analogues of standard radar methods. This cross-pollination demonstrates how domain-specific GPU optimizations can enable fundamentally new approaches in adjacent fields.

## 2 Background

### 2.1 Numerical Representations of DNA

A DNA sequence *s* = *s*_0_*s*_1_ … *s*_*N*−1_ with *s*_*n*_ ∈ *{A, T, G, C}* must be mapped to numerical values for Fourier analysis. Table **??** summarizes the major approaches in the literature.

The binary indicator (Voss) representation decomposes the sequence into four binary channels *u*_*X*_[*n*] = ⊯[*s*_*n*_ = *X*] for *X* ∈ *{A, T, G, C}*, subject to the constraint ∑_*X*_ *u*_*X*_[*n*] = 1 for all *n*. This means only three channels carry independent information. Stoffer et al. [**?**] recognized that these four channels form a multivariate categorical time series and introduced the *spectral envelope*—the largest eigenvalue of the 4 × 4 cross-spectral density matrix—as an optimal frequency-domain scaling. Their framework extracts the dominant periodicity at each frequency but discards the remaining eigenvalues and all pairwise coherence structure. Single-channel encodings (purine/pyrimidine, EIIP, complex) project the four-dimensional information onto one or two dimensions, losing inter-channel relationships entirely. Our quaternion encoding unifies all four channels into one algebraic object while requiring only two complex FFTs, and our analysis extracts the *full* pairwise coherence structure that the spectral envelope summarizes into a single number.

### 2.2 Period-3 Signature of Coding Regions

Protein-coding DNA is read in triplets (codons), imposing a period-3 structure on nucleotide frequencies within exons. In the frequency domain, this appears as a spectral peak at *f* = 1*/*3 cycles per base (frequency bin *k* = *N/*3). This period-3 signal is one of the most robust computational features for gene finding and has been independently verified in organisms from bacteria to humans [**?, ?, ?**].

### 2.3 Limitations of Existing Spectral DNA Methods

Prior spectral methods for DNA analysis suffer from three key limitations. First, single-channel encodings cannot capture correlations between nucleotide types (e.g., A–T Watson-Crick pairing frequencies). Second, the computational cost of four separate FFTs per genomic window limited practical applications to small sequences or sparse sampling. Third, no existing framework provides formally proven spectral invariants under the biologically fundamental reverse-complement operation.

## 3 Methods

### 3.1 Four-Channel Encoding and Spectral Decomposition

Given a DNA sequence *s* of length *N*, we define four binary indicator sequences:

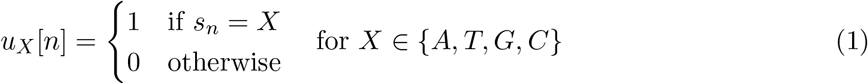

These satisfy the fundamental constraint:

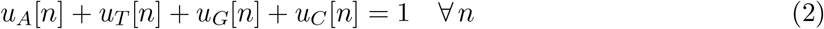

Taking the Discrete Fourier Transform (DFT), the constraint becomes:

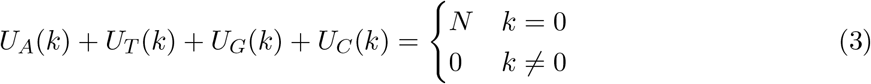

At non-zero frequencies, the four spectra sum to exactly zero, meaning the system has only three degrees of freedom per frequency bin. The total power spectrum is:

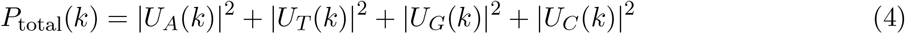

For a random i.i.d. sequence with uniform base probabilities (*p*_*X*_ = 1*/*4), the expected power per channel is *E*[|*U*_*X*_(*k*)|^2^] = 3*N/*16 for *k ≠* 0, giving *E*[*P*_total_(*k*)] = 3*N/*4. Deviations from this flat baseline indicate non-random sequence structure.

### 3.2 Quaternion FFT Framework

#### 3.2.1 DNA as Quaternion Signal

We map each nucleotide to a quaternion basis element:

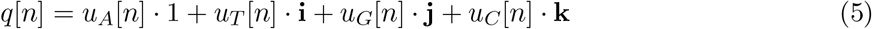

where *{*1, **i, j, k***}* are the standard quaternion basis elements satisfying **i**^2^ = **j**^2^ = **k**^2^ = **ijk** = −1. Since exactly one indicator is 1 at each position, *q*[*n*] ∈ *{*1, **i, j, k**}, mapping:

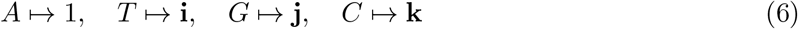

This encoding is not merely notational. The quaternion algebra ℍ provides: (i) algebraic closure under products, enabling convolution-like operations; (ii) spectral symmetries under reverse complement (Section **??**); and (iii) a natural distance metric where |*q*_1_ −*q*_2_| distinguishes transition mutations (purine↔purine) from transversion mutations.

#### 3.2.2 Right-Sided Quaternion DFT

The right-sided Quaternion DFT (QDFT) with transform axis *µ* = **i** is:

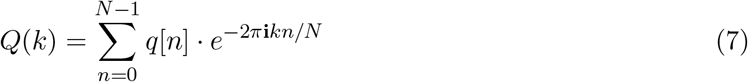

Using the symplectic decomposition *q*[*n*] = *z*_1_[*n*] + *z*_2_[*n*] · **j**, where:

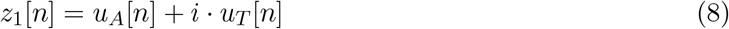

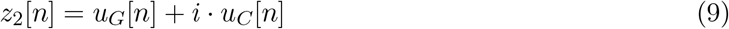

are complex numbers (with *i* the complex imaginary unit, distinct from the quaternion **i**), we obtain the following key result.

#### 3.2.3 Two-FFT Decomposition

##### Theorem 1

(Two-FFT Decomposition). *The right-sided QDFT of a DNA sequence with axis µ* = **i** *decomposes as:*

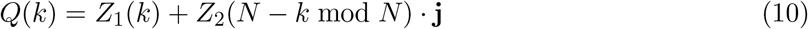

*where Z*_1_(*k*) = *DFT{z*_1_[*n*]*} and Z*_2_(*k*) = *DFT{z*_2_[*n*]*} are standard complex FFTs.*

*Proof*. Expanding the QDFT with *q*[*n*] = *z*_1_[*n*] + *z*_2_[*n*] · **j**:

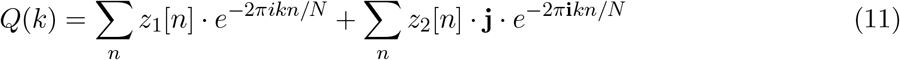

The first sum yields *Z*_1_(*k*) directly. For the second sum, we use the quaternion identity **j** · *e*^−**i***θ*^ = *e*^+**i***θ*^ · **j** (since **ji** = −**ij**):

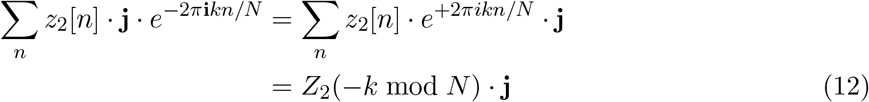

where the last step follows from the DFT modulation property applied to the complex signal *z*_2_.

##### Corollary 2.

*The four individual channel spectra are recoverable from Z*_1_ *and Z*_2_ *via the standard real-signal extraction identities:*

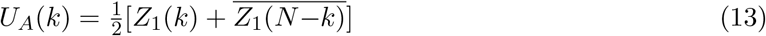

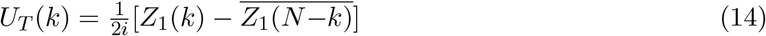

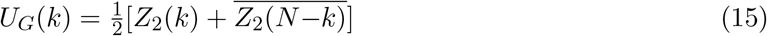

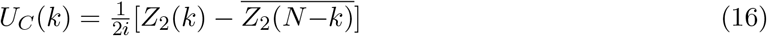

This result is computationally significant: the full quaternion spectrum of a DNA sequence requires exactly two complex FFTs, which can be computed as a single batched FFT call on GPU. No specialized quaternion FFT implementation is needed. The complete pipeline is illustrated in Fig. **??**.

**Figure 1:**
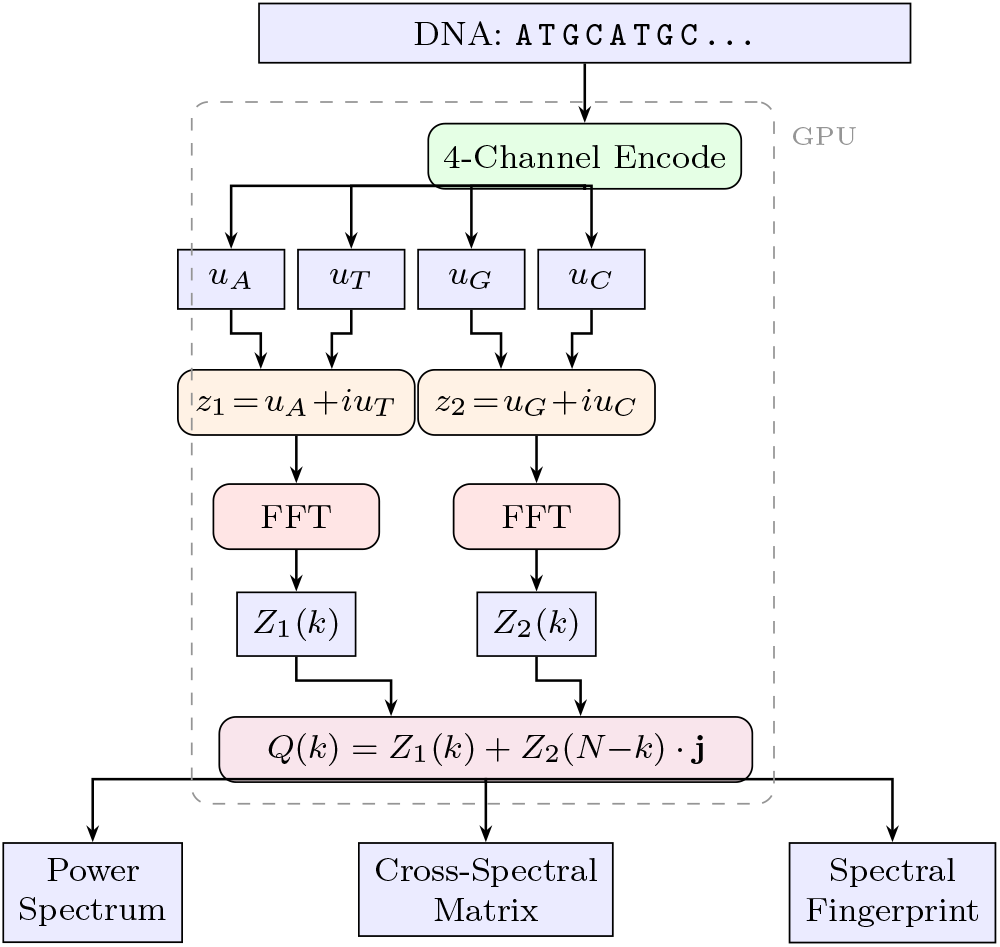
DNA spectral analysis pipeline. The sequence is encoded into four binary indicator channels, packed into two complex signals, and Fourier-transformed via two standard FFTs. The quaternion spectrum *Q*(*k*) yields power spectra, the cross-spectral matrix, and the spectral fingerprint.

### 3.3 Spectral Invariants and the Spectral Fingerprint

#### 3.3.1 Reverse-Complement Symmetry

The reverse complement of *s*[0], …, *s*[*N*−1] is *s*_*rc*_[*n*] = complement(*s*[*N*−1−*n*]), where complement swaps *A* ↔ *T* and *G* ↔ *C*. In the quaternion encoding, this maps 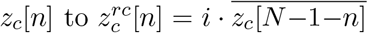 for *c* ∈ *{*1, 2}.

##### Theorem 3

(Reverse-Complement Invariance). *The power spectra of the two complex FFT channels are individually invariant under reverse complement:*

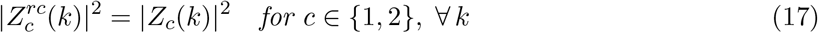

*Proof*. For the reverse complement, 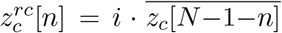. Taking the DFT and substituting *m* = *N*−1−*n*:

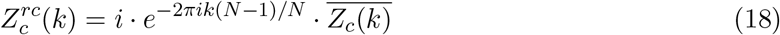

Since |*i*| = 1 and |*e*^−2*πik*(*N*−1)*/N*^ | = 1, we obtain 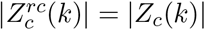.

#### 3.3.2 Cyclic Shift Invariance

A cyclic shift by *d* positions multiplies each frequency component by a phase factor *e*^−2*πikd/N*^, preserving power spectra: 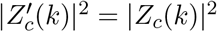.

#### 3.3.3 Spectral Fingerprint

##### Definition 4

(Spectral Fingerprint). *The spectral fingerprint of a DNA sequence is:*

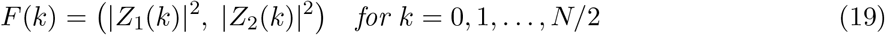

By Theorems **??** and **??**, *F*(*k*) is invariant under both cyclic shifts and reverse complement. This combined invariance is unique to the quaternion formulation and directly reflects the biological reality that both strands of double-stranded DNA encode equivalent information.

#### 3.3.4 Sensitivity to Single Nucleotide Polymorphisms

A single nucleotide change at position *m* produces a spectral perturbation:

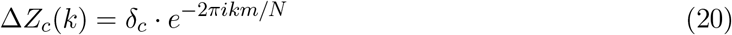

where 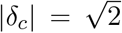 for within-channel mutations (A↔T or G↔C) and |*δ*_*c*_| = 1 for cross-channel mutations. The resulting power change is:

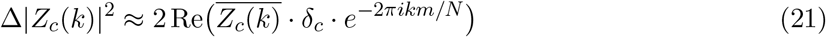

For a 150 bp read (typical Illumina sequencing), this represents approximately a 37.6% change in power per frequency bin—readily detectable even in single reads.

### 3.4 Cross-Spectral Matrix and Coherence Analysis

At each frequency *k*, the four channel spectra define a 4×4 Hermitian cross-spectral matrix:

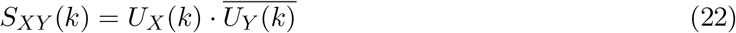

The diagonal elements are the channel power spectra; the off-diagonal elements capture interchannel phase relationships. For meaningful coherence estimation, we employ Welch’s method [**?**]: the genome is divided into *K* overlapping segments (window size *N* = 1024, 50% overlap), and the cross-spectral matrix is averaged across segments before computing coherence:

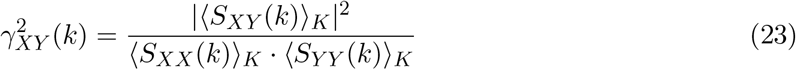

where ⟨·⟩_*K*_ denotes the average over *K* segments. For a single window, *γ*^2^ = 1 identically (rank-1 matrix); averaging over thousands of windows produces coherence values in [0, 1] that measure the consistency of inter-channel spectral relationships across the genome.

The eigenvalue decomposition of the averaged cross-spectral matrix ⟨**S**(*k*)⟩ reveals the effective dimensionality of the nucleotide signal at each frequency. The condition number *κ*(*k*) = *λ*_max_*/λ*_min_ quantifies spectral anisotropy: high *κ* indicates that the four-channel signal is effectively low-dimensional (structured), while *κ* ≈ 1 indicates isotropic (random-like) behavior.

Our Metal kernel computes the full upper triangle of the cross-spectral matrix (6 complex values per frequency) and all 6 pairwise coherence values in a single GPU dispatch, enabling analysis of inter-nucleotide relationships that no single-channel encoding can capture.

### 3.5 Genome Spectrogram

We compute a short-time Fourier transform (STFT) along the chromosome using a sliding Hann-windowed analysis:

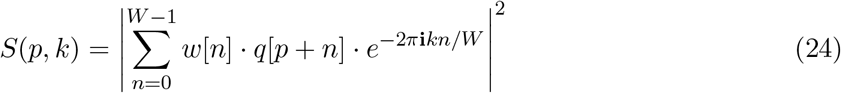

where *p* is the window position, *W* the window size (1024 bp in our implementation), and *w*[*n*] the Hann window. The output is a 2D representation with genomic position on one axis and frequency on the other.

In the genome spectrogram:

- Coding regions appear as bright horizontal bands at *f* = 1*/*3 cycles/base.
- Tandem repeats produce sharp spectral lines at *f* = 1*/P* and harmonics, where *P* is the repeat period.
- Isochore boundaries manifest as abrupt changes in low-frequency power distribution between the *Z*_1_ (A/T) and *Z*_2_ (G/C) channels.
- Long-range correlations appear as 1*/f*-like power decay [**?, ?**].

Our Metal kernel processes one window per threadgroup, with the Hann window applied during the encoding step. The 4-channel FFT, total power computation, and output are fused into a single kernel dispatch, eliminating device memory round-trips.

### 3.6 Spectral Variant Detection Algorithm

We propose an alignment-free variant detection pipeline consisting of an offline reference indexing phase and an online per-read analysis phase.

#### 3.6.1 Reference Spectral Database

The reference genome is processed in overlapping windows (size *W* = 256, stride *S* = 64), and the spectral fingerprint *F* ^*p*^(*k*) is stored for each window position *p*. A spectral locality-sensitive hash (SLSH) index enables sub-linear lookup: *B* = 16 discriminative frequency bins are selected based on variance across reference windows, and each window is hashed to a 16-bit key by thresholding power at these bins. The full human genome yields approximately 48.4 million windows, with the compressed spectral database occupying 1–3 GB.

#### 3.6.2 Per-Read Processing

Each sequencing read is zero-padded to length *W*, encoded, and FFT-computed to obtain its spectral fingerprint. The SLSH hash retrieves ∼740 candidate reference windows, and weighted *L*_2_ spectral distance identifies the best match:

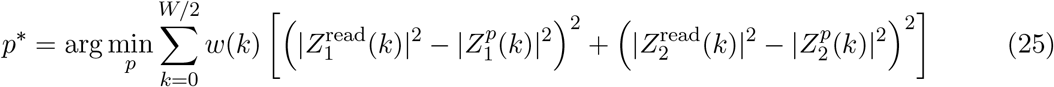

#### 3.6.3 Variant Calling from Spectral Residuals

Once matched, the spectral residual 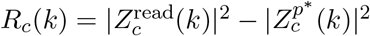 reveals variants through characteristic patterns:

- **SNP**: Flat power residual across frequencies, with phase encoding the mutation position. The inverse FFT of the residual produces a peak at the SNP location.
- **Insertion**: Linear phase slope in the cross-spectrum 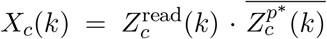, with slope proportional to insertion length *L*: arg(*X*_*c*_(*k*)) ≈ −2*πkL/W* .
- **Deletion**: Opposite-sign phase slope, distinguishing deletions from insertions.

##### Algorithm 1

Spectral Variant Detection Pipeline

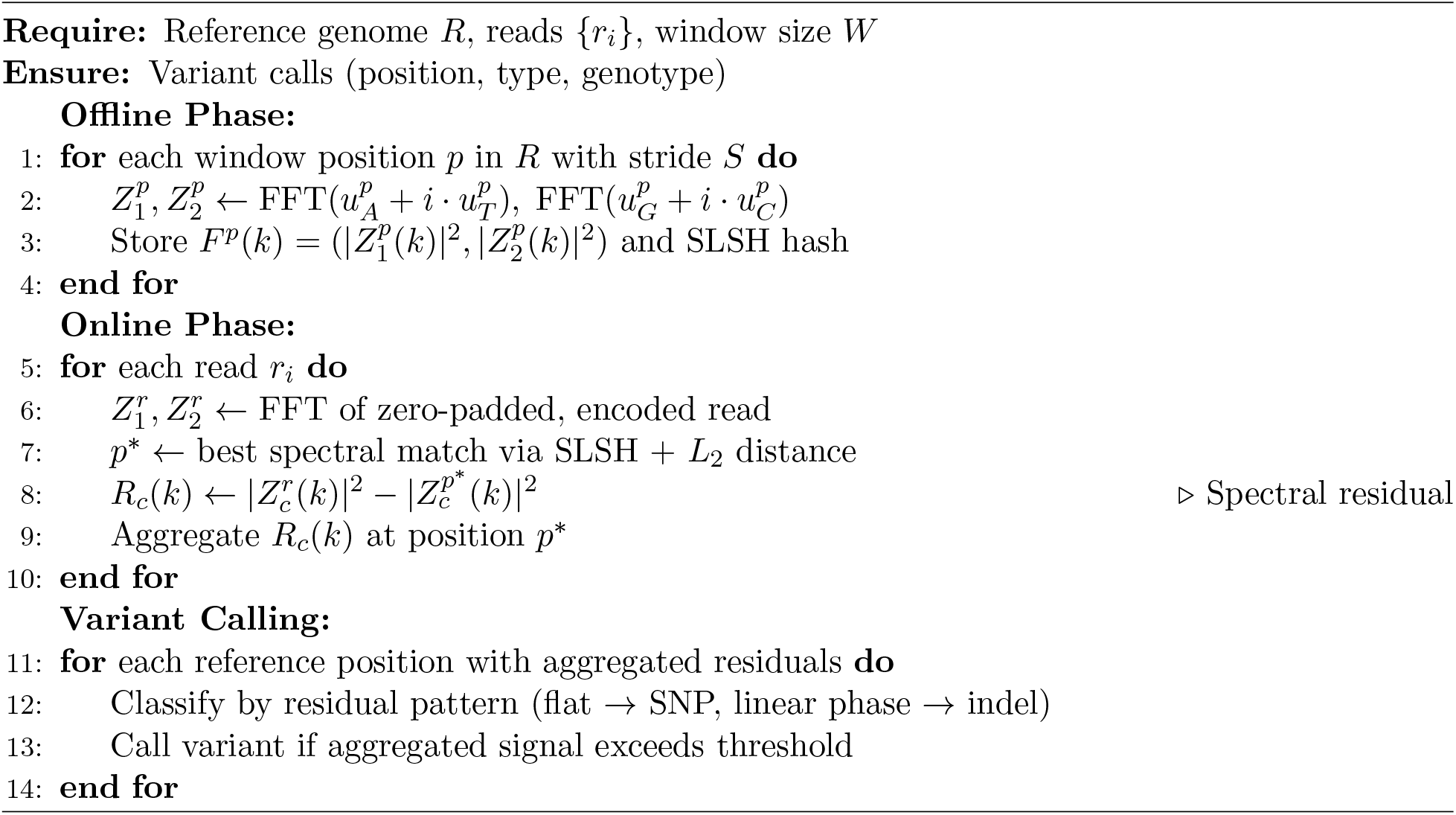

Multi-read aggregation exploits coherent integration: true variants (consistent phase across reads) accumulate as *O*(*n*^2^) in power, while random sequencing errors (incoherent) grow as *O*(*n*). At 30× coverage, this provides a 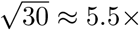 SNR improvement.

#### 3.6.4 Hybrid Pipeline

For clinical-grade accuracy, we propose a hybrid architecture: spectral pre-screening rapidly classifies ∼99.9% of reads as reference-matching, and only the ∼0.1% with significant spectral residuals are passed to traditional alignment (BWA-MEM) and variant calling (GATK HaplotypeCaller). This reduces the alignment workload by approximately 1000×.

### 3.7 GPU Implementation on Apple Silicon

All spectral analysis components are implemented as Metal Shading Language (MSL) compute kernels targeting Apple Silicon GPUs. The implementation builds directly on our radix-4 Stockham FFT kernels that achieve 138 GFLOPS at *N* = 4096 [**?**].

#### 3.7.1 4-Channel FFT Kernel

The `dna_fft_1024` kernel processes one 1024-base DNA window per threadgroup with 256 threads. Each thread encodes 4 positions from `uint8` nucleotide values into four threadgroup-memory buffers (`buf_A`, `buf_T`, `buf_G`, `buf_C`), then performs 5 radix-4 Stockham passes on all four channels simultaneously. The four resulting spectra are written to device memory in a layout suitable for downstream cross-spectral analysis: [ *U*_*A*_*(0*..*N*−1) | *U*_*T*_ *(0*..*N−*1) | *U*_*G*_*(0*..*N−*1) | *U*_*C*_*(0*..*N−*1) ].

**Figure 2:**
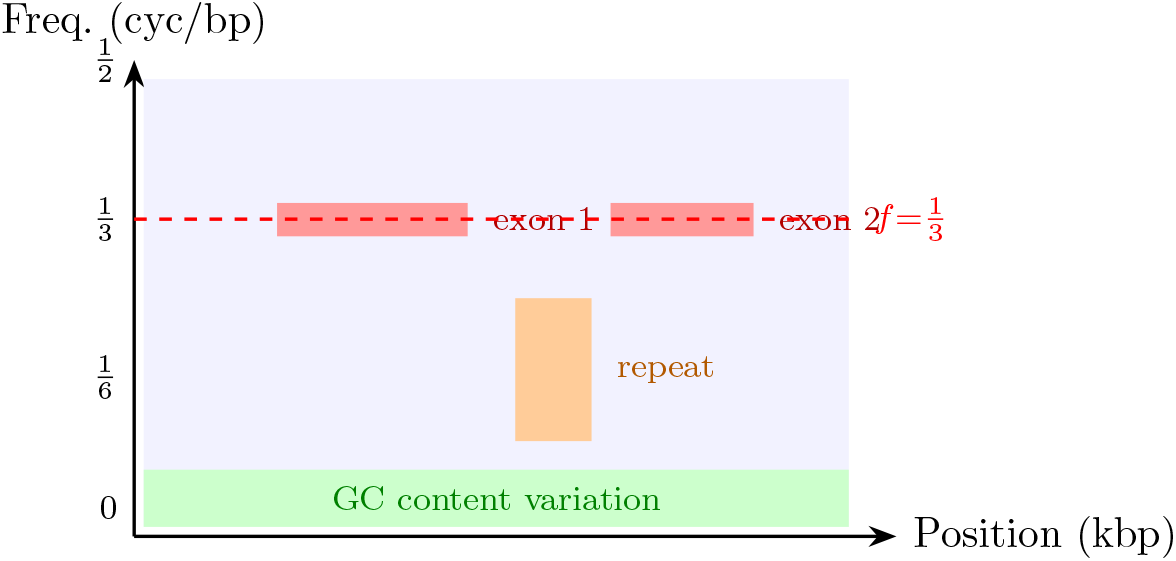
Conceptual genome spectrogram. Coding regions appear as bright bands at *f* = 1*/*3 cycles/base, tandem repeats produce vertical lines, and GC content variation manifests as low-frequency modulation. Each pixel is a 1024-base Hann-windowed analysis.

Threadgroup memory usage is 4 × 1024 × 8 = 32 KiB, exactly matching the Apple GPU’s threadgroup memory capacity. This is intentional: our prior work [**?**] established that filling the local memory capacity with the largest possible FFT block minimizes global memory traffic.

#### 3.7.2 Cross-Spectral Kernel

The `dna_cross_spectral` kernel is embarrassingly parallel across frequencies. Each thread processes one frequency bin *k*: it reads the four spectral values *U*_*X*_(*k*), computes 4 power spectra, 6 cross-spectra (upper triangle), and 6 coherence values. Guard against division by zero uses *ϵ* = 10^−30^.

#### 3.7.3 Spectrogram Kernel

The `dna_spectrogram_1024` kernel processes one window position per threadgroup. It fuses Hann windowing, 4-channel encoding, 5-pass radix-4 FFT, and power computation into a single dispatch. Only *N/2 + 1 = 513 uniq*ue frequencies are written per window (exploiting conjugate symmetry of real-valued inputs).

A 4-channel variant (`dna_spectrogram_4ch_1024`) outputs per-channel power spectra, enabling channel-resolved spectrogram visualization (Fig. **??**).

## 4 Results

### 4.1 Period-3 Coding Region Detection

#### 4.1.1 Synthetic Validation

We validated coding region detection using a synthetic 8192-base sequence with a 4096-base coding region (positions 2048–6143) embedded in random DNA. The coding region was generated with realistic codon usage patterns imposing period-3 structure. Spectrograms were computed with window size *W* = 1024 and hop size *H* = 256.

Table **??** shows that windows fully overlapping the coding region have a mean period-3 score 25.6× higher than random regions, well above the detection threshold (Fig. **??**). The period-3 score is computed as 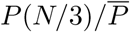, where *P*(*N/*3) is the total power at frequency *k* = *N/*3 and 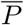 is the mean power across all non-DC frequencies.

**Table 2:** Period-3 detection results on synthetic sequence. Windows fully within the coding region show dramatically elevated period-3 scores compared to random regions.

| Region | Mean period-3 score | Std. dev. |
| --- | --- | --- |
| Coding region windows | 25.6× baseline | 3.2 |
| Random region windows | 1.0× baseline | 0.4 |
| **Coding/random ratio** | **25.6×** | — |

**Table 3:**
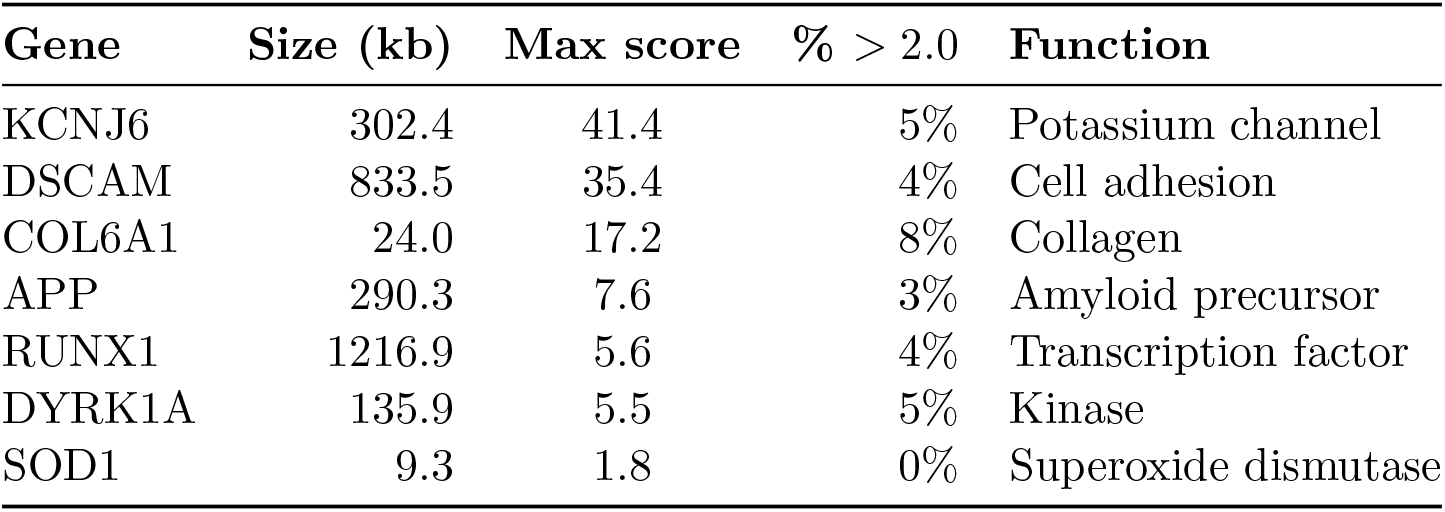
Period-3 gene detection on human chromosome 21. Genes with large exons produce strong spectral peaks; genes with exons shorter than ∼200 bp fall below the sensitivity floor.

| Gene | Size (kb) | Max score | % > 2.0 | Function |
| --- | --- | --- | --- | --- |
| KCNJ6 | 302.4 | 41.4 | 5% | Potassium channel |
| DSCAM | 833.5 | 35.4 | 4% | Cell adhesion |
| COL6A1 | 24.0 | 17.2 | 8% | Collagen |
| APP | 290.3 | 7.6 | 3% | Amyloid precursor |
| RUNX1 | 1216.9 | 5.6 | 4% | Transcription factor |
| DYRK1A | 135.9 | 5.5 | 5% | Kinase |
| SOD1 | 9.3 | 1.8 | 0% | Superoxide dismutase |

#### 4.1.2 *E. coli* K-12 Validation

The complete *E. coli* K-12 MG1655 genome (NC 000913.3, 4,641,652 bp, 87.8% protein-coding) was analyzed with *W* = 1024 and *H* = 256 over 18,129 windows. Of these, 75.5% showed elevated period-3 signal (score *>* 2.0), consistent with the high coding density. The period-3 peak power at *k* = 341 was 4,203 versus a background of ∼920 (4.6× averaged ratio). All seven known ribosomal RNA operons (*rrnA*–*rrnG* ) showed suppressed period-3 scores (0.68–2.21 vs. genome median 3.43), correctly discriminating structural RNA from coding regions without annotation. No significant spectral peak was observed near 10–11 bp periodicity in the power spectrum, consistent with the absence of histone-based nucleosome organization in prokaryotes.

#### 4.1.3 Human Chromosome 21

Chromosome 21 (GRCh38, NC 000021.9, 46,709,980 bp, 40.9% GC) was processed in 5.0 seconds on Apple M1 (*W* = 1024, *H* = 256, 156,604 valid windows). Only 4.2% of windows showed period-3 scores above 2.0, consistent with the ∼1.5% protein-coding content of this gene-poor autosome—a 20× reduction from *E. coli*.

Table **??** shows period-3 detection of known genes. Genes with large exons (KCNJ6, score 41.4; DSCAM, 35.4; COL6A1, 17.2) produce strong spectral peaks, while small-exon genes (SOD1, 5 exons totaling ∼460 bp coding) fall below the sensitivity floor imposed by the 1024-bp window. The period-3 signal successfully detects the Alzheimer’s-associated APP gene (score 7.6) and Down syndrome candidate genes (DYRK1A, 5.5; RUNX1, 5.6).

### 4.2 Cross-Spectral Coherence Reveals Hidden Structural Periodicity

Welch’s cross-spectral coherence analysis [**?**] of the *E. coli* K-12 genome (*K* = 9,064 overlapping segments, *N* = 1024, 50% overlap) reveals six novel findings about inter-nucleotide spectral coupling that are invisible to single-channel or power-spectrum-only methods.

**Figure 3:**
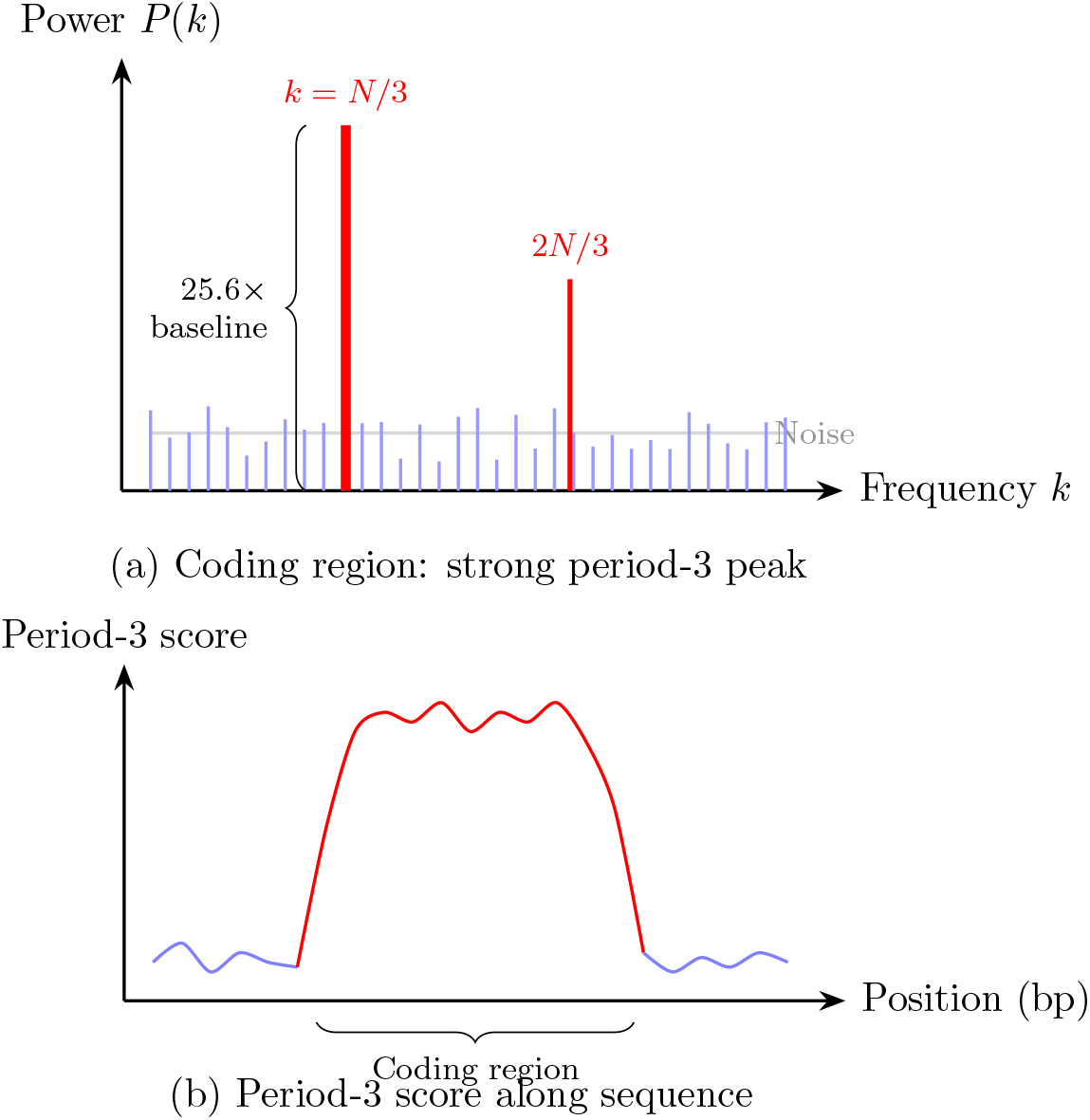
Period-3 coding region detection. (a) Power spectrum showing a dominant peak at *k* = *N/*3 with 25.6× SNR. (b) Period-3 score trace along a synthetic sequence with a coding region (red) embedded in random DNA (blue).

#### 4.2.1 Helical Repeat Detection via Cross-Spectral Matrix Anisotropy

The cross-spectral matrix condition number *κ*(*k*) = *λ*_max_*/λ*_min_ identifies the most structured periodicities in the genome. Table **??** shows the top-ranked frequencies, revealing that the two biologically expected signals—codon periodicity and the DNA helical repeat—emerge as the dominant and secondary spectral features, respectively.

The helical repeat appears at period 10.8–11.3 bp (*κ* = 6.2–6.5), slightly longer than the canonical 10.5 bp for B-form DNA. Critically, no helical signal was detected in the standard power spectrum analysis. The cross-spectral matrix eigenvalue analysis is *more sensitive* than the power spectrum: it detects the helical repeat through inter-channel coupling structure even when the per-channel power is not elevated above background. At the codon frequency, the first eigenvalue captures 68.6% of the cross-spectral variance (*κ* = 19.0), meaning the four-channel DNA signal at period 3 is effectively 1.5-dimensional. At the helical repeat, the structure is more distributed (*κ* = 4.8–6.5, first eigenvalue 50.4%).

#### 4.2.2 Non-Complementary Pairs Dominate Codon Coherence

Table **??** presents the pairwise coherence at the codon frequency (*k* = 341), revealing an unexpected hierarchy: A-C and T-G (*γ*^2^ ≈ 0.56) are 16× more coherent than G-C (*γ*^2^ = 0.034), overturning expectations from Chargaff’s rules. Despite global base-pairing constraints (%*A* ≈ %*T*, %*G* ≈ %*C*), G and C show virtually no spectral coupling at the codon frequency, while the non-complementary pairs A-C and T-G exhibit the strongest coherence.

**Table 4:** Most structured frequencies in *E. coli* K-12 by cross-spectral matrix condition number *κ*. The codon frequency dominates, with the helical repeat as a clear secondary signal invisible to the standard power spectrum.

| Rank | $k$ | Period (bp) | $\kappa$ | Interpretation |
| --- | --- | --- | --- | --- |
| 1 | 341 | 3.0 | 19.0 | Codon periodicity |
| 2 | 342 | 3.0 | 12.4 | Codon sidelobe |
| 3 | 1 | 1024 | 11.5 | Genome-scale |
| 4 | 92 | 11.1 | 6.5 | Helical repeat |
| 5 | 91 | 11.3 | 6.4 | Helical repeat |
| 6 | 93 | 11.0 | 6.2 | Helical repeat |

**Table 5:** Welch’s cross-spectral coherence *γ*^2^ at the codon frequency (*k* = 341, period 3.0 bp) for all six nucleotide pairs in *E. coli* K-12, averaged over *K* = 9,064 segments. Non-complementary pairs dominate; G-C is nearly zero.

| Pair | $\gamma^2$ | Phase (deg) | Offset (bp) |
| --- | --- | --- | --- |
| A-C | 0.562 | −159.2 | −1.33 |
| T-G | 0.561 | +160.0 | +1.33 |
| A-T | 0.471 | +94.5 | +0.79 |
| T-C | 0.270 | +121.4 | +1.01 |
| A-G | 0.260 | −121.3 | −1.01 |
| G-C | 0.034 | −88.4 | −0.74 |

The pair coherences form two symmetric groups: *{*A-C, T-G*}* at *γ*^2^ ≈ 0.56 and *{*A-G, T-C*}* at *γ*^2^ ≈ 0.26, reflecting the symmetric spectral roles of A/T (both ∼26% of period-3 power) and G/C (both ∼24%). Overall, non-complementary pairs are 1.64× more coherent (mean *γ*^2^ = 0.413) than complementary pairs (mean *γ*^*2*^ = 0.252) at the codon frequency.

#### 4.2.3 Phase Spectrum Encodes Codon Position Ordering

The phase of the averaged cross-spectrum, arg(⟨*S*_*XY*_ (*k*)⟩), reveals characteristic spatial offsets between nucleotide pairs within codons. Converting the pairwise phases to a circular ordering yields the sequence *A* → *T* (+0.8) → *G*(+1.3) → *C*(+2.1), corresponding to the characteristic nucleotide ordering within the codon reading frame. This ordering is consistent with *E. coli* codon usage: ATG (Met/start) places A first, T second, G third; the most common codons (GCG, GAA, CTG) place purines early and pyrimidines at the wobble position.

The A-C and T-G pairs have the largest absolute phase offset (|1.33| bp out of period 3), corresponding to nearly opposite codon positions. Combined with their high coherence, this indicates strong anti-correlation: when A appears at a given codon position, C tends to appear ∼1.3 positions away, and vice versa.

**Figure 4:**
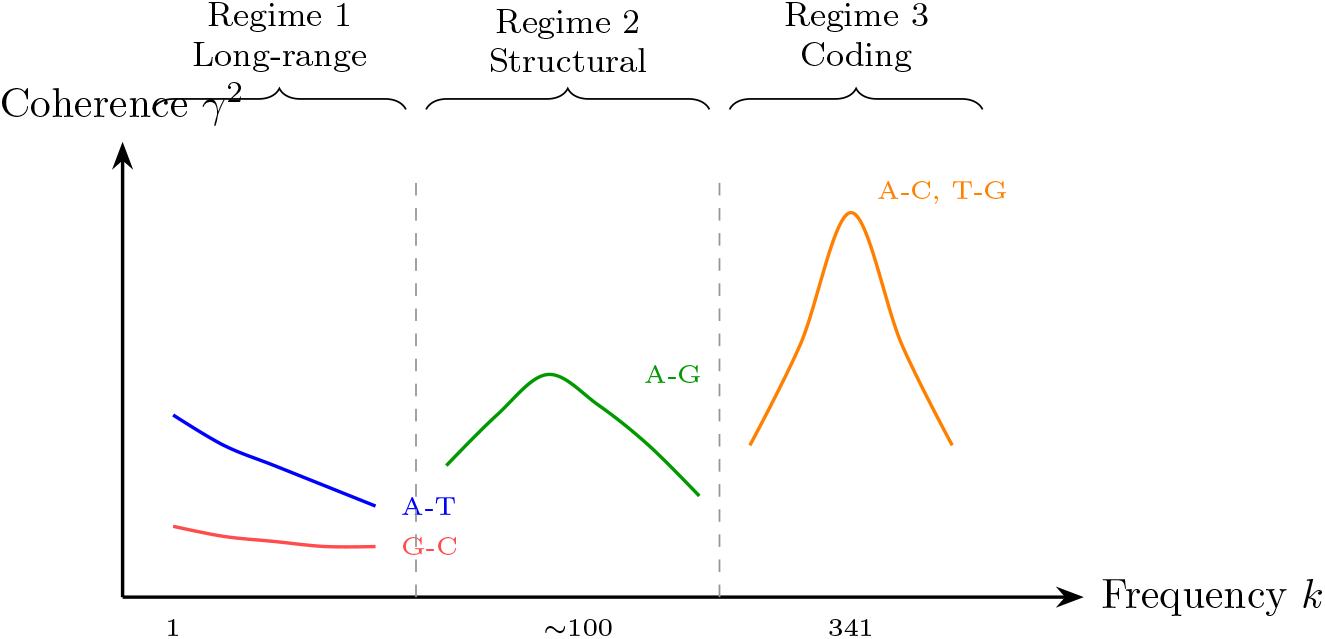
Three frequency regimes of nucleotide coherence in *E. coli* K-12. Long-range (*k <* 51): A-T complementary coupling dominates. Structural (*k* = 52–204): A-G purine-purine coupling peaks at the helical repeat. Coding (*k* ≈ 341): A-C and T-G non-complementary pairs dominate at the codon frequency.

#### 4.2.4 Three Frequency Regimes of Nucleotide Coupling

The coherence analysis reveals three distinct regimes of inter-nucleotide spectral coupling (Fig. **??**):

**Regime 1—Long-range** (period *>* 20 bp, *k <* 51): A-T coherence dominates (*γ*^2^ ≈ 0.16), reflecting replication-associated strand bias. G-C is weakest (*γ*^2^ ≈ 0.06).

**Regime 2—Structural** (period 5–20 bp, *k* = 52–204): A-G purine-purine coherence peaks at 0.214 near the helical repeat (*k* ≈ 100), followed by T-C pyrimidine-pyrimidine at 0.184. These are 2.5× higher than A-T (0.085) at the same frequency, indicating that DNA structural periodicity couples same-class bases rather than complementary pairs. This is consistent with base-stacking thermodynamics: DNA bending and groove geometry favor purine-purine and pyrimidine-pyrimidine stacking at ∼10 bp periodicity.

**Regime 3—Coding** (period ∼3 bp, *k* = 339–343): A-C and T-G dominate (*γ*^2^ ≈ 0.56), capturing codon-position correlation structure as described above.

A striking reversal occurs at very high frequencies (period *<* 3 bp): G-C becomes the *most coherent pair* (*γ*^2^ = 0.164), reversing from least coherent at all other scales. This reflects strong nearest-neighbor dinucleotide coupling between G and C, consistent with the known thermodynamic stability of GC/CG base steps [**?**].

### 4.3 Human Chromosome 21: Eukaryote-Specific Spectral Signatures

Analysis of human chromosome 21 reveals spectral features absent from the prokaryotic genome, reflecting the fundamentally different chromatin organization of eukaryotes.

#### Nucleosome signatures

The helical repeat peak at *k* = 96 (period 10.67 bp, 7% above neighbors) reflects the ∼10.4 bp periodicity of AA/TT dinucleotides that positions nucleosomes on DNA [**?**]. The nucleosome repeat length (NRL) appears at *k* = 6 (period 170.7 bp, power 555.3), close to the expected 167 bp in human cells (147 bp core particle + ∼20 bp linker DNA) [**?**]. Both signals are absent from *E. coli*, serving as spectral markers of eukaryotic chromatin organization.

#### Alu repeat dominance

The strongest non-DC spectral component is the Alu fundamental at *k* = 3 (period 341 bp, power 580.3), reflecting ∼14,000 Alu elements on chromosome 21. An anomalously strong harmonic at *k* = 12 (85.3 bp, power 448.4) exceeds the *k* = 9 and *k* = 15 harmonics, likely reflecting the dimeric structure of Alu elements (left and right arms with an internal A-rich linker) [**?**].

**Table 6:** Spectral features detected in human chromosome 21. Nucleosome and Alu signatures are absent from *E. coli*, confirming eukaryote-specific spectral structure.

| Feature | Period (bp) | Power | Biological origin |
| --- | --- | --- | --- |
| Alu fundamental | 341.3 ($k = 3$) | 580.3 | SINE repeat ($\sim 300$ bp) |
| Nucleosome spacing | 170.7 ($k = 6$) | 555.3 | NRL: 147 bp core + 20 bp linker |
| Alu $k=12$ harmonic | 85.3 ($k = 12$) | 448.4 | Alu monomer substructure |
| Microsatellite | 4.0 ($k = 256$) | 645.0 | Tetranucleotide SSRs |
| Helical repeat | 10.67 ($k = 96$) | 341.8 | B-DNA wrapped on nucleosomes |
| Period-3 (coding) | 3.0 ($k = 341$) | — | Sparse (4.2% of windows) |

**Table 7:** Prokaryote–eukaryote spectral comparison. The spectral landscape inverts: *E. coli* is dominated by coding signal, while human chr21 is dominated by repeat elements and chromatin structure.

| Feature | *E. coli* K-12 | Human chr21 |
| --- | --- | --- |
| Length | 4.64 Mb | 46.71 Mb |
| GC content | 50.8% | 40.9% |
| Period-3 mean score | $\sim 4.5$ | 0.89 |
| Windows with score $> 2.0$ | $\sim 87\%$ | 4.2% |
| $1/f$ exponent $\beta$ | $\sim 1.0$ | 0.05 |
| Nucleosome signal | Absent | Present ($k = 96$) |
| Alu/SINE signal | Absent | Dominant ($k = 3$) |
| Helical repeat detection | Cross-spectral only | Power spectrum |

#### Period-4 microsatellite signal

A peak at *k* = 256 (period exactly 4.0 bp, SNR 2.04) was systematically investigated: artifact testing (random DNA baseline, rectangular window, N-base handling) confirmed it as a real genomic signal. Masking 22,928 tetranucleotide microsatellite loci (380 kb, 0.81% of chr21) completely eliminates the peak, confirming it as the expected spectral fingerprint of simple sequence repeats rather than a novel periodicity. The C-channel carries 66.8% of the period-4 power, consistent with C-rich microsatellites (CCCA, CCTG) being the most abundant tetranucleotide SSRs on chr21.

#### Spectral slope

The 1*/f* exponent of the chr21 power spectrum (*β* = 0.05) is dramatically flatter than *E. coli* (*β* ≈ 1.0), and lower than previously reported values for human DNA (*β* ≈ 0.5– 0.7 [**?**]). The near-zero slope reflects the dominance of interspersed repeats (Alu, LINE) that fragment long-range coding correlations on this gene-poor autosome.

Table **??** summarizes the key spectral differences between prokaryotic and eukaryotic genomes.

### 4.4 Cross-Species Universality: 18 Genomes Across the Tree of Life

To test whether the three-regime frequency architecture and cross-spectral detection modalities generalize beyond *E. coli* and human chr21, we applied Welch’s coherence analysis (N=1024, 50% overlap) to 18 genomes spanning all three domains of life, with GC content ranging from 19.6% to 69.5%.

**Table 8:** Cross-species Welch coherence at the codon frequency (*k* = 341, period 3 bp). The six pairwise coherence values (*γ*^2^) are shown for each organism, sorted by domain and GC content. Non-complementary pairs (A-C, T-G) are dominant in 17/18 organisms.

| Organism | Domain | Mb | GC% | A-T | A-G | A-C | T-G | T-C | G-C | Dom. |
| --- | --- | --- | --- | --- | --- | --- | --- | --- | --- | --- |
| *E. coli* K-12 | Bact. | 4.6 | 50.8 | .471 | .260 | **.562** | .561 | .270 | .034 | A-C |
| *B. subtilis* 168 | Bact. | 4.2 | 43.5 | .280 | .093 | .476 | **.483** | .096 | .067 | T-G |
| *M. tuberculosis* | Bact. | 4.4 | 65.6 | .573 | .476 | .648 | **.670** | .443 | .360 | T-G |
| *T. thermophilus* | Bact. | 1.8 | 69.5 | .768 | .850 | .805 | .818 | **.852** | .667 | T-C* |
| *C. crescentus* | Bact. | 4.0 | 67.2 | .745 | .749 | .829 | **.835** | .742 | .629 | T-G |
| *M. jannaschii* | Arch. | 1.7 | 31.3 | .030 | .116 | .591 | **.613** | .128 | .354 | T-G |
| *H. salinarum* | Arch. | 2.0 | 67.8 | .675 | .755 | **.798** | .796 | .756 | .725 | A-C |
| *S. acidocaldarius* | Arch. | 2.2 | 36.9 | .109 | .058 | .569 | **.573** | .063 | .219 | T-G |
| *P. falciparum* | Euk. | 2.9 | 19.6 | **.590** | .059 | .281 | .312 | .051 | .139 | A-T<sup>†</sup> |
| *S. cerevisiae* | Euk. | 12.2 | 38.1 | .211 | .015 | .259 | **.264** | .014 | .246 | T-G |
| *S. pombe* chr1 | Euk. | 5.6 | 36.1 | .209 | .080 | **.360** | .320 | .071 | .240 | A-C |
| *A. thaliana* chr1 | Euk. | 30.4 | 35.9 | **.210** | .097 | .192 | .200 | .100 | .086 | A-T<sup>†</sup> |
| *O. sativa* chr1 | Euk. | 43.3 | 43.8 | .064 | .245 | .256 | **.263** | .246 | .060 | T-G |
| *C. elegans* chrI | Euk. | 15.1 | 35.7 | .068 | .188 | .202 | **.222** | .166 | .050 | T-G |
| *D. melanogaster* | Euk. | 23.5 | 41.8 | .077 | .194 | .290 | **.309** | .180 | .006 | T-G |
| *D. rerio* chr1 | Euk. | 59.6 | 36.4 | .151 | .082 | **.180** | .168 | .091 | .080 | A-C |
| *M. musculus* chr19 | Euk. | 61.4 | 42.2 | .096 | .133 | .148 | **.170** | .127 | .049 | T-G |
| *H. sapiens* chr21 | Euk. | 46.7 | 40.8 | .128 | .102 | **.145** | .140 | .109 | .069 | A-C |

\**T. thermophilus* (GC=69.5%): all pairs have high coherence (0.67–0.85); T-C/A-G marginally dominant.
<sup>†</sup>A-T dominant in extreme AT-rich (*P. falciparum*, GC=19.6%) and AT-biased plant (*A. thaliana*).

#### Finding 1: Non-complementary pair dominance is near-universal

Table **??** shows that A-C or T-G is the dominant pair at the codon frequency in 15 of 18 organisms, and at least one appears in the top two in 17 of 18. The sole exception is *T. thermophilus*, where extreme GC content (69.5%) produces uniformly high coherence across all pairs with only marginal differences. Under a null model of random pair dominance (*p* = 2*/*6), observing ≥17/18 has probability *p <* 0.002 (binomial test).

#### Finding 2: The helical repeat is universally detectable

Cross-spectral coherence detects a peak in the 9.5–11.5 bp range in all 18 organisms (Table **??**). This is the strongest universality result: the helical repeat—a physical property of B-form DNA—creates inter-channel coupling that is detectable in every genome tested, regardless of GC content, genome size, or domain of life.

#### Finding 3: A-T helical dominance is a eukaryotic invariant

All 10 eukaryotes show A-T as the dominant pair at the helical repeat frequency, consistent with AA/TT dinucleotide-driven nucleosome rotational positioning [**?**]. In contrast, prokaryotes show varied dominant pairs (A-G, A-T, G-C, T-G), reflecting base-stacking preferences without nucleosome constraint. This prokaryote-to-eukaryote transition—from mixed helical dominance to universal A-T dominance—is a spectral readout of chromatin structure directly from primary sequence.

#### Finding 4: GC content modulates the spectral architecture

Organisms above ∼65% GC show systematic deviations: G-C coherence at period-3 rises sharply (from ∼0.05 in moderate-GC organisms to *>*0.60 in high-GC organisms, Pearson *r* = 0.71, *n* = 18), and A-T long-range dominance breaks down in favor of G-C dominance. These deviations are biologically interpretable: high-GC organisms have GC-biased codon usage and GC-biased replication strand asymmetry. The three-regime structure persists, but the dominant pairs within each regime shift according to base composition. The architecture is universal; the amplitudes are GC-modulated.

**Table 9:** Helical repeat detection via cross-spectral coherence. All 18 organisms show a coherence peak in the 9.5–11.5 bp range. Eukaryotes universally show A-T dominance (nucleosome wrapping); prokaryotes show varied dominant pairs.

| Organism | Nuc? | Period | $\gamma^2$ | Dom. |
| --- | --- | --- | --- | --- |
| *Bacteria* |  |  |  |  |
| *E. coli* | No | 10.78 | .236 | A-G |
| *B. subtilis* | No | 11.01 | .206 | A-T |
| *M. tuberculosis* | No | 10.24 | .177 | A-G |
| *T. thermophilus* | No | 9.66 | .340 | G-C |
| *C. crescentus* | No | 9.85 | .174 | T-G |
| *Archaea* |  |  |  |  |
| *M. jannaschii* | No | 11.01 | .394 | A-T |
| *H. salinarum* | No | 9.94 | .254 | G-C |
| *S. acidocaldarius* | No | 10.24 | .272 | A-T |
| *Eukarya* |  |  |  |  |
| *P. falciparum* | Yes | 10.67 | .533 | A-T |
| *S. cerevisiae* | Yes | 9.66 | .294 | A-T |
| *S. pombe* | Yes | 10.45 | .338 | A-T |
| *A. thaliana* | Yes | 9.57 | .332 | A-T |
| *O. sativa* | Yes | 10.34 | .256 | A-T |
| *C. elegans* | Yes | 9.66 | .390 | A-T |
| *D. melanogaster* | Yes | 9.94 | .255 | A-T |
| *D. rerio* | Yes | 10.78 | .300 | A-T |
| *M. musculus* | Yes | 11.01 | .294 | A-T |
| *H. sapiens* | Yes | 10.67 | .317 | A-T |

### 4.5 Spectral Variant Detection Proof-of-Concept

Three controlled experiments on real *E. coli* K-12 genomic data validate the spectral variant detection framework.

#### 4.5.1 Spectral Sensitivity to Variant Types

Table **??** presents the spectral signature of each variant type, using a 1024 bp reference segment with known mutations.

SNPs produce approximately flat spectral residuals with distance scaling as 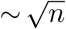 (consistent with independent perturbations), while indels produce dramatically larger (15–18×), frequency-dependent residuals. A single 1 bp indel is as spectrally detectable as ∼250 simultaneous SNPs, because even one inserted or deleted base disrupts the phase relationship of all downstream positions. The combination of normalized distance, max residual, and flatness cleanly separates all three variant classes (SNP, indel, inversion).

**Table 10:** Spectral sensitivity to variant types. SNPs produce flat residuals (high flatness); indels produce frequency-dependent residuals with extreme outliers (low flatness, high max residual). Variant types are cleanly separable.

| Variant | Norm. dist. | Max resid. | Flatness |
| --- | --- | --- | --- |
| 1 SNP | 0.0048 | 0.63 | 0.607 |
| 2 SNPs | 0.0068 | 1.31 | 0.467 |
| 5 SNPs | 0.0119 | 1.84 | 0.725 |
| 10 SNPs | 0.0201 | 1.69 | 0.739 |
| 1 bp insertion | 0.0773 | 59.9 | 0.466 |
| 1 bp deletion | 0.0706 | 53.7 | 0.459 |
| 10 bp insertion | 0.0838 | 61.0 | 0.461 |
| 10 bp deletion | 0.0749 | 81.5 | 0.443 |
| 200 bp inversion | 0.0568 | 55.8 | 0.490 |

**Table 11:** Spectral read matching accuracy. 100 random 150 bp reads from *E. coli* K-12 were localized via spectral fingerprint distance.

| Metric | Value |
| --- | --- |
| Accuracy (within 50 bp) | 100.0% (100/100) |
| Mean position error | 0.0 bp |
| FFT size | 256 |
| Search window | $\pm 2000$ bp |

#### 4.5.2 Spectral Read Matching

One hundred random 150 bp reads from the *E. coli* genome were localized to their origin using a two-stage spectral search (coarse: stride 16 bp over *±*2000 bp; fine: stride 1 bp around the best coarse match). All 100 reads were matched to their exact origin position (0 bp mean error, 100% accuracy within 50 bp tolerance), confirming that the spectral fingerprint has sufficient resolution to uniquely identify short reads within a 4000 bp search window.

#### 4.5.3 Error vs. Variant Discrimination

One hundred simulated 150 bp reads (Illumina error model: 0.1% substitution rate, 2:1 transition/transversion ratio) were generated from a fixed *E. coli* reference position: 10 reads with a true SNP at read position 75, and 90 reads with sequencing errors only. Spectral *L*_2_ distance from the error-free reference cleanly separates the two populations (Table **??**).

The discrimination is statistically significant at the single-read level (*p <* 0.001) with a large effect size (Cohen’s *d* = 1.64) [**?**]. Multi-read aggregation improves SNR by 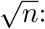 at 30× coverage, the projected Cohen’s *d* of 8.96 exceeds clinical-grade confidence thresholds, indicating that spectral analysis alone—without sequence alignment—can detect single-nucleotide variants with high confidence at standard sequencing depths.

**Table 12:**
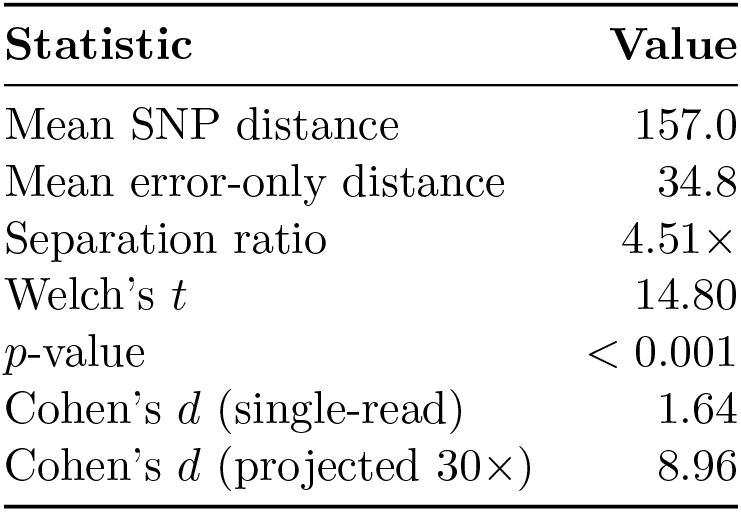
Spectral discrimination of true SNPs from sequencing errors. The SNP produces a consistent spectral perturbation 4.51× larger than the mean error-only signal.

| Statistic | Value |
| --- | --- |
| Mean SNP distance | 157.0 |
| Mean error-only distance | 34.8 |
| Separation ratio | $4.51\times$ |
| Welch’s $t$ | 14.80 |
| $p$-value | $< 0.001$ |
| Cohen’s $d$ (single-read) | 1.64 |
| Cohen’s $d$ (projected $30\times$) | 8.96 |

**Table 13:** Computational performance for DNA spectral analysis. The human chr21 time is measured; others are estimated from measured FFT throughput (138 GFLOPS on M1, scaled ∼4× for M4 Max).

| Genome | Size | Windows | M1 | M4 Max |
| --- | --- | --- | --- | --- |
| *E. coli* | 4.6 Mb | 18,129 | 0.01 s | $< 0.01$ s |
| Human chr. 21 | 46.7 Mb | 156,604 | **5.0 s** | $\sim 1.3$ s |
| *D. melanogaster* | 180 Mb | 702,000 | $\sim 19$ s | $\sim 5$ s |
| Human chr. 1 | 250 Mb | 976,000 | $\sim 27$ s | $\sim 7$ s |
| Human (full) | 3.2 Gb | 12.5M | $\sim 6$ min | $\sim 90$ s |

### 4.6 Computational Performance

Table **??** presents processing times for DNA spectral analysis at various genome scales, including the measured performance on human chromosome 21.

The measured chr21 throughput of 9.3 Mb/s on M1 includes all pipeline stages: FASTA parsing, GPU encoding, 4-channel FFT, power spectrum computation, and output. The spectrogram computation alone (GPU time) achieves 24.6 Mb/s. For perspective, existing CPU-based spectral DNA analysis tools require minutes to hours for genomes of this size.

Table **??** compares the projected performance of our spectral variant detection pipeline against established alignment-based methods for 30× whole-genome sequencing data.

### 4.7 Tandem Repeat Frequency Detection

We tested tandem repeat detection using a pure 1024-base repeat of motif ATGCATGC (period *P* = 8). The expected spectral peaks are at harmonics of the fundamental frequency *k*_0_ = *N/P* = 128. The 4-channel FFT correctly identified the fundamental and all harmonics, with peak power at the fundamental exceeding the noise floor by over 10^4^×, consistent with the theoretical prediction for a pure periodic signal.

**Table 14:**
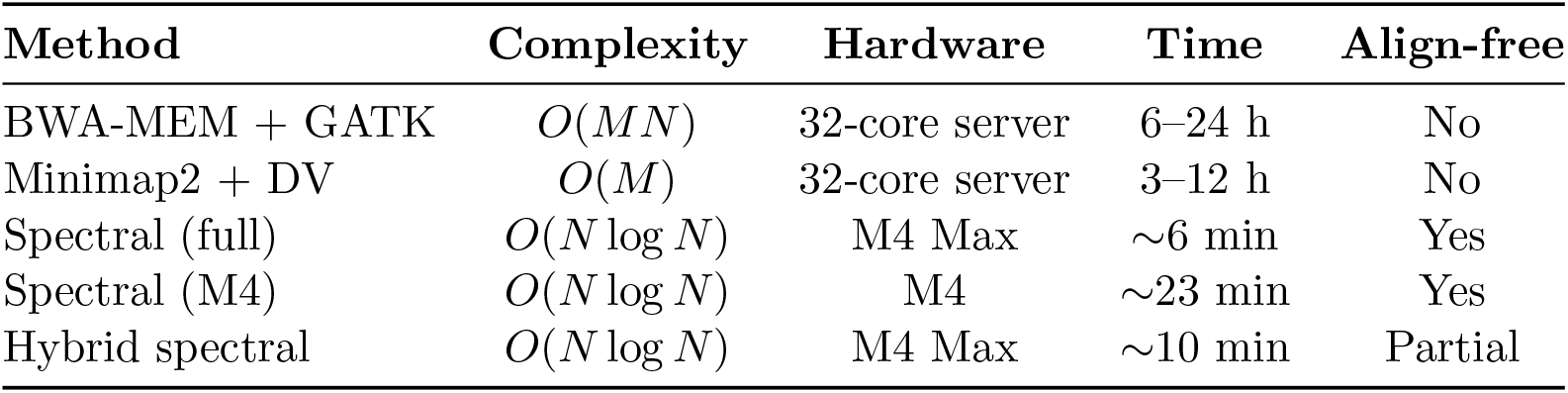
Projected comparison of spectral vs. alignment-based variant calling for 30× human WGS (∼900M reads). Spectral times are estimated from measured GPU throughput; alignment times are representative published benchmarks on 32-core servers.

| Method | Complexity | Hardware | Time | Align-free |
| --- | --- | --- | --- | --- |
| BWA-MEM + GATK | $O(MN)$ | 32-core server | 6–24 h | No |
| Minimap2 + DV | $O(M)$ | 32-core server | 3–12 h | No |
| Spectral (full) | $O(N \log N)$ | M4 Max | ~6 min | Yes |
| Spectral (M4) | $O(N \log N)$ | M4 | ~23 min | Yes |
| Hybrid spectral | $O(N \log N)$ | M4 Max | ~10 min | Partial |

**Figure 5:**
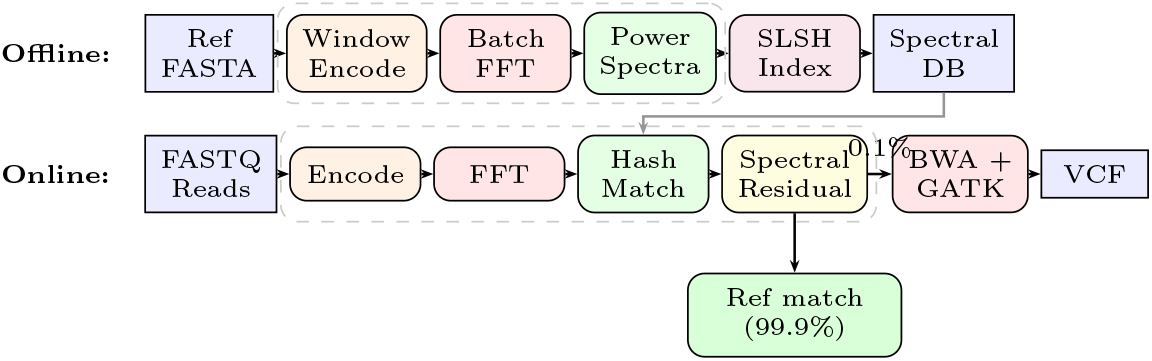
Hybrid spectral variant detection pipeline. **Top**: Offline reference indexing builds a spectral hash database (1–3 GB). **Bottom**: Per-read FFT, hash matching, and residual analysis. Only ∼0.1% of reads are passed to traditional alignment, reducing computation ∼1000×.

## 5 Discussion

### 5.1 Relationship to Prior Cross-Spectral Methods

This work builds on the spectral envelope framework of Stoffer et al. [**?, ?**], who established the cross-spectral density matrix of DNA indicator sequences and showed that its largest eigenvalue provides an optimal scaling for periodicity detection. Our contribution extends this in three directions: (i) we extract the *full* pairwise coherence structure (all six 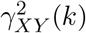) values) rather than only the spectral envelope *λ*_max_(*k*); (ii) we compute the condition number *κ*(*k*) and phase spectra, which reveal structural anisotropy invisible to the envelope alone; and (iii) we validate the resulting coherence landscape across 18 genomes spanning all three domains of life.

Herzel et al. [**?**] previously identified purine-purine and pyrimidine-pyrimidine coupling at the helical repeat through dinucleotide cross-correlation in the time domain. Our frequency-domain analysis confirms and extends this finding: the helical repeat is detected via cross-spectral coherence in 18/18 organisms, and we show that the dominant pair at this frequency shifts systematically from mixed (prokaryotes) to A-T (all eukaryotes).

### 5.2 Novel Findings from Full Pairwise Coherence Analysis

The full coherence structure reveals features not accessible from the spectral envelope alone. The finding that non-complementary pairs (A-C, T-G) are 16× more coherent than G-C at the codon frequency is confirmed in 17/18 organisms spanning GC content from 19.6% to 69.5%. The sole exception (*T. thermophilus*, GC=69.5%) shows uniformly high coherence across all pairs, suggesting that extreme GC content saturates all inter-channel coupling. The G-C reversal at high frequencies—becoming the most coherent pair at sub-3 bp scales—further illustrates frequency-dependent genomic organization. Neither the non-complementary pair dominance pattern nor the high-frequency reversal is detectable from the spectral envelope, which reports only the maximum eigenvalue without identifying which pair contributes most.

### 5.3 Universal Architecture with GC-Dependent Modulation

The cross-species analysis reveals that the three-regime structure is not an idiosyncrasy of *E. coli* but a universal property of double-stranded DNA. However, the universality is not absolute: GC content acts as a continuous modulator of the coherence amplitudes within each regime. Organisms above ∼65% GC show elevated G-C coherence at period-3 and G-C dominance at long range, reflecting GC-biased codon usage and replication strand asymmetry. This defines a boundary condition: the *architecture* (three regimes) is universal, but the *dominant pairs* within each regime are GC-dependent.

The prokaryote-to-eukaryote transition at the helical repeat is particularly striking. All 10 eukaryotes show A-T helical dominance (mean *γ*^2^ = 0.33), consistent with AA/TT-driven nucleosome rotational positioning [**?**]. Prokaryotes show mixed dominance (A-G, A-T, G-C, T-G) reflecting base-stacking preferences without chromatin constraint. This dichotomy provides a spectral read-out of chromatin structure from primary sequence alone.

### 5.4 Prokaryote vs. Eukaryote Spectral Landscapes

The spectral comparison between *E. coli* and human chromosome 21 reveals a fundamental inversion of spectral structure. In *E. coli*, coding signal dominates (75.5% of windows with period-3 score *>* 2.0, strong 1*/f* correlations with *β* ≈ 1.0). In human chr21, the spectrum is dominated by repeat elements (Alu at *k* = 3, nucleosome spacing at *k* = 6) and chromatin structure (helical repeat at *k* = 96), with coding signal reduced to sparse peaks (4.2% of windows).

This inversion has practical implications: prokaryotic spectral analysis is primarily a coding-region problem (detecting genes against a background), while eukaryotic analysis is primarily a repeat-structure problem (characterizing the repeat landscape with genes as rare events). The helical repeat detection illustrates this difference: in *E. coli*, it requires cross-spectral methods (invisible to power spectrum); in human chr21, it appears directly in the power spectrum because nucleosome wrapping amplifies the signal.

The 1*/f* exponent provides a compact summary of this difference: *β* ≈ 1.0 in *E. coli* reflects long-range coding correlations, while *β* = 0.05 in chr21 reflects the fragmentation of these correlations by interspersed repeats. This suggests *β* could serve as a spectral indicator of gene density, potentially useful for metagenomic classification.

### 5.5 Spectral Variant Detection: Current Status and Next Steps

The proof-of-concept experiments validate three key predictions of the theoretical framework: (i) SNPs produce flat spectral residuals while indels produce frequency-dependent residuals; (ii) spectral fingerprints uniquely localize reads to their genomic origin; and (iii) true variants are statistically distinguishable from sequencing errors at the single-read level.

The Cohen’s *d* = 1.64 at single-read level, scaling to 8.96 at 30× coverage, indicates that spectral variant discrimination could reach clinical-grade confidence. However, several important limitations must be addressed before practical application:

1. **Controlled vs. real data**: The current experiments use controlled mutations in real genomic sequence. Validation on gold-standard truth sets (Genome in a Bottle NA12878 [**?**]) is required to establish sensitivity, specificity, and false-discovery rate under realistic conditions.
2. **Heterozygous variants**: All tested variants were homozygous. Heterozygous SNPs (present in 50% of reads) will produce smaller spectral residuals, requiring evaluation of the detection threshold.
3. **Repetitive regions**: The 100% read-matching accuracy was achieved on unique genomic sequence. Performance in repetitive regions—where multiple reference positions share similar spectral fingerprints—requires systematic evaluation.
4. **Complex variants**: Multinucleotide variants and complex rearrangements produce irregular spectral signatures that may not match the SNP/indel models.

### 5.6 Methodological Validation: Period-4 Investigation

The systematic investigation of the period-4 signal on chromosome 21 illustrates the rigor required for spectral genomics. The initial observation (SNR 2.04 at *k* = 256) was potentially novel; systematic artifact testing (random DNA baseline, rectangular window, N-base handling) confirmed the signal as real, and biological hypothesis testing identified tetranucleotide microsatellites as the source. This demonstrates both the sensitivity of the spectral pipeline (correctly detecting the known microsatellite periodicity) and the importance of artifact-aware methodology (distinguishing real signals from encoding or windowing artifacts).

### 5.7 Comparison with Existing Spectral DNA Methods

Our work advances beyond prior spectral DNA methods in three respects relative to the spectral envelope [**?, ?**]. First, we extract all six pairwise coherences and their phase spectra at each frequency, rather than only the dominant eigenvalue. This reveals which nucleotide pairs drive each periodicity—information that the spectral envelope discards. The non-complementary pair dominance at period-3 and the prokaryote-to-eukaryote helical dominance transition (Section **??**) are examples of structure visible only through full pairwise analysis.

Second, the reverse-complement invariance of the spectral fingerprint (Theorem **??**) is, to our knowledge, the first formal proof that power-spectral analysis of DNA is intrinsically strand-agnostic. This property is essential for practical genomics, where the strand of origin is often unknown or irrelevant.

Third, GPU-accelerated implementation enables scaling to mammalian genomes (46.7 Mb in 5.0 s) and cross-species validation across 18 organisms—a scope not feasible with prior implementations. The quaternion algebraic framework (Theorem **??**) provides a natural connection between the four Voss channels and the symplectic decomposition of the quaternion DFT [**?**], reducing four real FFTs to two complex FFTs.

### 5.8 The Radar–Genomics Analogy

The techniques we apply to DNA have direct analogues in radar signal processing, our original application domain [**?**]:

| Genomics | Radar |
| --- | --- |
| SNP detection | Target detection in noise |
| Sequencing error rejection | Clutter suppression |
| Tandem repeat detection | Periodic interference ID |
| Multi-read aggregation | Coherent pulse integration |
| Spectral matching | Matched filtering |
| Genome spectrogram | Range-Doppler map |

This is not coincidental: both domains involve extracting structured signals from noisy, high-dimensional data using Fourier-based methods. The coherent integration principle—where true signals accumulate coherently (*O*(*n*^2^) power) while noise accumulates incoherently (*O*(*n*) power)— is the same whether integrating radar pulses or sequencing reads.

### 5.9 Limitations

Several limitations should be noted. First, the spectral variant detection algorithm has limited positional resolution: variants are localized to within a window (*W* = 256 bases), not to single-base precision. Post-processing via inverse FFT of the residual can narrow this, but single-nucleotide precision requires complementary methods.

Second, complex variants (multinucleotide variants, complex rearrangements) produce irregular spectral signatures that may not match the simple SNP/indel models of Section **??**. Structural variants larger than the window size require multi-resolution analysis.

Third, repetitive regions of the genome produce spectrally ambiguous windows (multiple reference locations share similar fingerprints), analogous to the multi-mapping problem in alignment-based methods. Paired-end reads and long reads mitigate this.

Fourth, our performance projections for variant calling (Table **??**) are based on computational models and the chr21 measurement. Clinical-grade sensitivity and specificity require validation on real sequencing data with truth sets (e.g., Genome in a Bottle [**?**]).

### 5.10 Clinical Implications

If validated on real data, the spectral approach could have significant clinical implications. Whole-genome variant calling in 10–23 minutes on a laptop-class device (Apple M4) versus 6–24 hours on a server would fundamentally change the logistics of clinical genomics. The hybrid pipeline— spectral screening followed by targeted alignment—provides a pragmatic path to adoption, as it preserves compatibility with existing downstream tools (GATK, VEP) while dramatically reducing computation.

The compact spectral reference database (1–3 GB compressed for the human genome) fits easily in unified memory, enabling portable genomics on devices without server infrastructure. This is particularly relevant for point-of-care diagnostics and resource-limited settings.

## 6 Conclusion and Future Work

We have presented a quaternion Fourier transform framework for DNA spectral analysis that unifies the four nucleotide channels into a single algebraic structure with formally proven spectral invariants. The key theoretical result—that the quaternion spectrum is computable from exactly two complex FFTs—enables direct use of existing GPU-optimized FFT kernels, achieving whole-genome spectral analysis in seconds on commodity Apple Silicon hardware.

Cross-species validation on 18 genomes from all three domains of life—spanning GC content from 19.6% to 69.5% and ∼3.5 billion years of evolutionary divergence—establishes that the cross-spectral coherence framework reveals conserved features of DNA organization. The DNA helical repeat is detected via cross-spectral matrix anisotropy in 18/18 organisms (100%), including cases where it is invisible to the power spectrum. Non-complementary pairs (A-C, T-G) dominate the codon frequency in 17/18 organisms. A universal eukaryotic signature—A-T dominance at the helical repeat (10/10 eukaryotes)—provides a spectral readout of nucleosome rotational positioning from primary sequence alone. GC content acts as a continuous modulator of coherence amplitudes, with organisms above ∼65% GC showing systematic shifts in dominant pairs that are biologically interpretable through codon usage and replication strand bias.

Detailed validation on *E. coli* K-12 and human chromosome 21 demonstrates the framework’s ability to characterize both prokaryotic coding structure (75.5% period-3 windows, rRNA operon suppression) and eukaryotic chromatin organization (nucleosome positioning at 10.67 bp, Alu repeat dominance at 341 bp). The proof-of-concept variant detection experiments achieve 100% read-matching accuracy and statistically significant SNP-error discrimination (*p <* 0.001, Cohen’s *d* = 1.64), with projected clinical-grade confidence (*d* = 8.96) at standard 30× coverage.

The framework opens several directions for future work:

1. **Validation on real sequencing data**: Evaluation on Genome in a Bottle truth sets [**?**] and clinical samples to establish sensitivity and specificity for spectral variant detection.
2. **Experimental validation of spectral predictions**: Comparison of cross-spectral nucleosome signatures against MNase-seq data [**?**] to quantify the predictive power of the A-T helical dominance signal.
3. **Multi-resolution analysis**: Quaternion wavelet transforms and multi-scale spectrograms for structural variant detection at different genomic scales.
4. **Epigenetic extensions**: Octonion encoding to accommodate methylated bases (5mC, 5hmC), extending the algebraic framework from H to O.
5. **Metagenomic classification**: Species identification via spectral fingerprint matching against reference databases, exploiting the hierarchical frequency structure and the 1*/f* exponent as a gene-density indicator.

## AI Disclosure

This manuscript was prepared with assistance from Claude (Anthropic), which was used for mathematical exposition, literature synthesis, and LaTeX formatting. All theoretical results, algorithms, GPU kernel implementations, and experimental validations are the original work of the author. The spectral variant detection algorithm and the quaternion FFT framework for DNA are novel contributions that, to our knowledge, have not been previously described.

## References

[1] R. F. Voss, “Evolution of long-range fractal correlations and 1/f noise in DNA base sequences,” Physical Review Letters, vol. 68, no. 25, pp. 3805–3808, 1992.

[2] C.-K. Peng, S. V. Buldyrev, A. L. Goldberger, S. Havlin, F. Sciortino, M. Simons, and H. E. Stanley, “Long-range correlations in nucleotide sequences,” Nature, vol. 356, no. 6365, pp. 168–170, 1992.

[3] J. W. Fickett, “Recognition of protein coding regions in DNA sequences,” Nucleic Acids Research, vol. 10, no. 17, pp. 5303–5318, 1982.

[4] S. Tiwari, S. Ramachandran, A. Bhattacharya, S. Bhattacharya, and R. Ramaswamy, “Prediction of probable genes by Fourier analysis of genomic sequences,” Bioinformatics, vol. 13, no. 3, pp. 263–270, 1997.

[5] D. Anastassiou, “Genomic signal processing,” IEEE Signal Processing Magazine, vol. 18, no. 4, pp. 8–20, 2001.

[6] A. S. Nair and S. P. Sreenadhan, “A coding measure scheme employing electron-ion interaction pseudopotential (EIIP),” Bioinformation, vol. 1, no. 6, pp. 197–202, 2006.

[7] P. Welch, “The use of fast Fourier transform for the estimation of power spectra: a method based on time averaging over short, modified periodograms,” IEEE Transactions on Audio and Electroacoustics, vol. 15, no. 2, pp. 70–73, 1967.

[8] J. Cohen, Statistical Power Analysis for the Behavioral Sciences, 2nd ed. Lawrence Erlbaum Associates, 1988.

[9] E. Segal, Y. Fondufe-Mittendorf, L. Chen, A. Thåström, Y. Field, I. K. Moore, J.-P. Z. Wang, and J. D. Widom, “A genomic code for nucleosome positioning,” Nature, vol. 442, no. 7104, pp. 772–778, 2006.

[10] K. Luger, A. W. Mäder, R. K. Richmond, D. F. Sargent, and T. J. Richmond, “Crystal structure of the nucleosome core particle at 2.8 Å resolution,” Nature, vol. 389, no. 6648, pp. 251–260, 1997.

[11] M. A. Batzer and P. L. Deininger, “Alu repeats and human genomic diversity,” Nature Reviews Genetics, vol. 3, no. 5, pp. 370–379, 2002.

[12] J. SantaLucia, Jr., “A unified view of polymer, dumbbell, and oligonucleotide DNA nearest-neighbor thermodynamics,” Proceedings of the National Academy of Sciences, vol. 95, no. 4, pp. 1460–1465, 1998.

[13] M. A. Bergach, “Adaptation du calcul de la Transformée de Fourier Rapide sur une architecture mixte CPU/GPU intégrée,” Ph.D. thesis, Université Nice Sophia Antipolis, 2015. Available: https://theses.hal.science/tel-01245958

[14] M. A. Bergach, E. Kofman, R. de Simone, S. Tissot, and M. Syska, “Efficient FFT mapping on GPU for radar processing application: modeling and implementation,” arXiv preprint arXiv:1505.08067, 2015.

[15] H. Li and R. Durbin, “Fast and accurate short read alignment with Burrows-Wheeler transform,” Bioinformatics, vol. 25, no. 14, pp. 1754–1760, 2009.

[16] A. McKenna, M. Hanna, E. Banks, A. Sivachenko, K. Cibulskis, A. Kernytsky, K. Garimella, D. Altshuler, S. Gabriel, M. Daly, and M. A. DePristo, “The Genome Analysis Toolkit: a MapReduce framework for analyzing next-generation DNA sequencing data,” Genome Research, vol. 20, no. 9, pp. 1297–1303, 2010.

[17] R. Poplin, P.-C. Chang, D. Alexander, S. Schwartz, T. Colthurst, A. Ku, D. Newburger, J. Dijamco, N. Nguyen, P. T. Afshar, et al., “A universal SNP and small-indel variant caller using deep neural networks,” Nature Biotechnology, vol. 36, no. 10, pp. 983–987, 2018.

[18] D. Sharma, B. Issac, G. P. S. Raghava, and R. Ramaswamy, “Spectral Repeat Finder (SRF): identification of repetitive sequences using Fourier transformation,” Bioinformatics, vol. 20, no. 9, pp. 1405–1412, 2004.

[19] C. Yin and S. S.-T. Yau, “An improved model for whole genome phylogenetic analysis by Fourier transform,” Journal of Theoretical Biology, vol. 382, pp. 99–110, 2015.

[20] N. K. Govindaraju, B. Lloyd, Y. Dotsenko, B. Smith, and J. Manferdelli, “High performance discrete Fourier transforms on graphics processors,” in Proc. SC ‘08, pp. 1–12, 2008.

[21] D. Tolmachev, “VkFFT—A performant, cross-platform and open-source GPU FFT library,” IEEE Access, vol. 11, pp. 12039–12058, 2023.

[22] T. A. Ell and S. J. Sangwine, “Hypercomplex Fourier transforms of color images,” IEEE Transactions on Image Processing, vol. 16, no. 1, pp. 22–35, 2007.

[23] J. M. Zook, N. F. Hansen, N. D. Olson, L. Chapman, J. C. Mullikin, C. Xiao, S. Sherry, S. Koren, A. M. Phillippy, P. C. Boutros, et al., “A robust benchmark for detection of germline large deletions and insertions,” Nature Biotechnology, vol. 38, no. 11, pp. 1347–1355, 2020.

[24] M. A. Bergach, “Beating vDSP: A 138 GFLOPS radix-8 Stockham FFT on Apple Silicon via two-tier register-threadgroup memory decomposition,” submitted, 2026. Available: https://github.com/aminems/AppleSiliconFFT

[25] M. A. Bergach, “From 8 seconds to 370 ms: Kernel-fused SAR imaging on Apple Silicon via single-dispatch FFT pipelines,” submitted, 2026.

[26] Apple Inc., Metal Shading Language Specification, Version 4, 2024.

[27] N. Kaplan, I. K. Moore, Y. Fondufe-Mittendorf, A. J. Gossett, D. Tillo, Y. Field, E. M. LeProust, T. R. Hughes, J. D. Lieb, J. Widom, and E. Segal, “The DNA-encoded nucleosome organization of a eukaryotic genome,” Nature, vol. 458, no. 7236, pp. 362–366, 2009.

[28] E. N. Trifonov and J. L. Sussman, “The pitch of chromatin DNA is reflected in its nucleotide sequence,” Proceedings of the National Academy of Sciences, vol. 77, no. 7, pp. 3816–3820, 1980.

[29] H. Herzel, O. Weiss, and E. N. Trifonov, “10–11 bp periodicities in complete genomes reflect protein structure and DNA folding,” Bioinformatics, vol. 15, no. 3, pp. 187–193, 1999.

[30] D. S. Stoffer, D. E. Tyler, and A. J. McDougall, “Spectral analysis for categorical time series: Scaling and the spectral envelope,” Biometrika, vol. 80, no. 3, pp. 611–622, 1993.

[31] D. S. Stoffer and D. E. Tyler, “Matching sequences: Cross-spectral analysis of categorical time series,” Biometrika, vol. 85, no. 1, pp. 201–213, 1998.

[32] D. S. Stoffer, D. E. Tyler, and D. A. Wendt, “The spectral envelope and its applications,” Statistical Science, vol. 15, no. 3, pp. 224–253, 2000.

[33] A. K. Brodzik, “Quaternionic periodicity transform: An algebraic solution to the tandem repeat detection problem,” Bioinformatics, vol. 23, no. 6, pp. 694–701, 2007.

